# Haplo-pheno association for *OsNRT1.1* paralog in rice reveals superior haplogroup with high nitrate uptake efficiency

**DOI:** 10.1101/2023.12.07.570681

**Authors:** Dharani Elangovan, Renu Pandey, Sandeep Sharma, Balaji Balamurugan, Neha Anand, Adhip Das, Tarun Kumar, Ranjith Kumar Ellur, Sanjay Kalia, Jagadish Rane

**Author notes:** Corresponding author Renu Pandey.

## Abstract

Haplotype-based breeding approaches hold promise for enhancing crop improvement strategies, allowing for targeted selection of superior genetic combinations to develop high-yielding and resilient varieties. The current study aimed at identification of *NRT1.1* nitrate transporter haplotype that could serve as “donors” in haplotype-based breeding. We phenotyped 272 rice accessions in hydroponics with sufficient and low nitrogen (N) for nitrate uptake efficiency. By employing principal component and hierarchical cluster analysis, the accessions were grouped into N efficient, intermediate, and inefficient clusters. Haplotype analysis unveiled the presence of two haplogroups for *OsNRT1.1A*, three for *OsNRT1.1B*, and five for *OsNRT1.1C*. Through haplo-pheno association, the comparison of mean trait values revealed H2 and H3 as the superior haplotypes (SH) for OsNRT1.1A and OsNRT1.1B, respectively. In the case of OsNRT1.1C, H3 and H1 emerged as SH within the N-efficient cluster. Conversely, the inferior haplotypes (IH) consisted of H1 in OsNRT1.1A, H3 in OsNRT1.1B, and H3 and H2 in OsNRT1.1C within the N-inefficient cluster. However, relative expression of *OsNRT1.1* (with specific paralogs) in contrasting rice accessions revealed that a few of the inferior accession exhibited higher expression levels in the root but lower in the shoot, which might have contributed to their N-inefficiency. Furthermore, amino acid change at position 403 (Isoleucine to Valine) in inferior accessions influences the active site OsNRT1.1C protein causing N-inefficiency. Ours is the first report on haplotype analysis of *NRT1.1* gene demonstrating its genetic diversity, as well as its association with phenotype will have potential implications for improving nitrate uptake efficiency.

## Introduction

Rice (*Oryza sativa*) is one of the most important cereal crops of the world, grown in a wide range of climatic zones. For nearly 80% of the dietary needs, more than 50% of the world’s population relies on rice as a staple food (Fukagawa and Ziska 2019). In the harvest year 2021–2022, about 509.87 million metric tonnes of rice were consumed worldwide. Furthermore, rice accounts for one-fifth of all calories consumed by humans globally, making it the most important food crop in terms of human nutrition and calorie intake. The seven-fold rise in the usage of nitrogen (N) fertilizers has been associated with the doubling of agricultural food output over the last four decades. The judicious and effective application of fertilizers can greatly increase production and enhance rice quality (Alam et al. 2009).

Nitrogen holds a pivotal position within plants, serving as a component in various biomolecules like proteins, nucleic acids, and chlorophyll, which are integral to essential physiological processes. In soil, the two primary inorganic forms of N are nitrate (NO_3_^-^) and ammonium (NH_4_^+^) (Li et al. 2017). In aerobic conditions, nitrate represents the primary form of N, while in flooded environments or acidic soils, ammonium typically prevails. The key role of N is promoting carbohydrate accumulation in rice culms, leaf sheaths, and grains during both the pre-heading and ripening stages (Swain et al. 2010). The key growth phases of rice crop, that is, the early vegetative stage and the panicle initiation stage, are influenced by N nutrition. It is found that the belowground parts were more susceptible to N throughout the vegetative growth, especially during emergence of panicles, while aboveground parts were mostly affected during the tillering period (Thi Nong et al. 2020). For transplanted rice, the application of 120 kg N ha^-1^ at crucial growth stages is suggested (Thind and Gupta 2010). In India, farmers tend to apply large amounts of N fertilizer in excess of plant requirements due to the substantial subsidy on urea fertilizer (Ladha et al. 2005). However, the major portion of N is lost from the soil through volatilization, denitrification, and leaching, which consequently lead to the pollution of both air and water and an increase in production cost (Plett et al. 2020). As the world’s demand for resources continues to rise, the primary focus of N management is to fulfil food security requirements while reducing the emission of unutilized reactive N species, such as NO_x_, which is significantly more harmful to the environment, being 300 times more potent than CO_2_. Inefficient utilization of N-based fertilizers by crops, resulting in N losses in various forms such as nitrous oxide, ammonia, and NO_x_ into the atmosphere, as well as nitrates/ammonium ions leaching into water bodies, presents substantial economic and environmental challenges (Abrol et al. 2012).

Plants adapt to low or high nitrate concentrations in soil *via* high-affinity and low-affinity nitrate-uptake systems, respectively (Feng et al. 2011). High-affinity transport systems (HATS) operate at low nutrient concentrations (K_m_∼µM). At higher amounts (K_m_∼mM), LATS are predominantly responsible for absorption allowing for a substantial uptake of substrate while maintaining high substrate availability (Tsay 2014; Parker et al. 2014). Two *NRT1* (low-affinity) transporters and four *NRT2* (high-affinity) transporters have been reported in root nitrate uptake in *Arabidopsis thaliana*: *NRT1.1, NRT1.2, NRT2.1, NRT2.2, NRT2.4*, and *NRT2.5* (Okamoto et al. 2003). However, in most growth environments, *NRT1.1* and *NRT2.1* play the most critical roles in root nitrate uptake, while the other four NRTs contribute just minimal (Ye et al. 2019). So far, in rice, more than 80 NRT1/PTR, 4 NRT2, and 2 NAR2 members have been identified but only a few members of the NRT1/PTR family are reported (Araki et al. 2006; Cai et al. 2008). The first low-affinity nitrate transporter identified in rice, *OsNRT1*, mediates nitrate uptake in roots (Li et al. 2017). In Arabidopsis, *NRT1.1* is a dual-affinity transporter, which switches affinity *via* phosphorylation and dephosphorylation at the threonine amino acid residue (designated as Thr_101_) (Tsay 2014).

The NRT1/PTR family of secondary active transporters belongs to the Major Facilitator Superfamily (MFS), which uses the proton electrochemical gradient to promote substrate absorption into the cell (Reddy et al. 2012). NRT1.1 belongs to the NITRATE TRANSPORTER 1/PEPTIDE TRANSPORTER FAMILY (NPF) that exists across all organisms. The NRT1.1 protein consists of a canonical MFS fold which is formed by 12 transmembrane (TM)-spanning alpha helices organized into two bundles, amino-terminal (TM1-TM6) and carboxyterminal (TM7-TM12). The Thr_101_ is located at the base of TM3 and directs attention to a hydrophobic pocket made of TM2 and TM4 residues. (Parker et al. 2014). Phosphorylation at Thr_101_ in AtNRT1.1 leads to the separation of dimers, potentially enhancing structural adaptability. This structural flexibility boost, in turn, elevates the transport rate, resulting in alterations in K_m_, primarily attributed to the enhancing nitrate transport rate (Tsay 2014). The dynamic nature of N as well as its susceptibility to loss from soil and plant systems creates a unique and demanding environment for optimal management. Changes in nutrient uptake and utilization patterns produced by nutritional stresses will eventually reflect changes in molecular activity.

To meet future food demand, harnessing the allelic diversity of genetic resources of various crops is crucial. It is always worthwhile to look for better alleles of a gene in order to create and maintain natural genetic diversity (Neelam et al. 2017). The recently developed and efficient ’haplotype assembly’ concept enables the integration of superior variants of genes that have been identified (Bevan et al. 2017). In this context, haplotype analysis has become a useful tool for investigating genetic variants and determining how certain sets of genetic markers are inherited together. A haplotype is a set of single nucleotide polymorphism (SNPs) on one chromosome that are inherited from one parent (Silverman 2007), whereas a haplogroup is a group of organisms/genotypes sharing the same haplotype from a common ancestor. A haplotype might be limited to a single gene or it can be bigger and consist of many genes. In recent years, haplotype-based genome-wide association studies (GWAS) have discovered significant QTLs and candidate genes for a number of traits in crops like rice, wheat, soybean, etc. (Contreras-Soto et al. 2017; Ogawa et al. 2021; Hamazaki and Iwata 2020). For example, the presence of multiple haplotypes for sucrose synthase gene reflects heterozygosity but the enhanced fixation of elite alleles in modern sugarcane germplasm may be limiting its further genetic advancements (Zhang et al. 2013). While haplotype analysis has been conducted on several genes for various traits, but the studies related to mineral nutrition in agricultural crops, particularly rice, remains relatively scarce. In the present study, we phenotyped 272 rice accessions to assess their ability to acquire nitrate efficiently during the seedling stage under varying N levels. Simultaneously, haplotype analysis of these accessions, derived from the 3K Rice Genome panel, was carried out to pinpoint the superior haplotypes for the paralogs of OsNRT1.1 that can be employed in haplotype-based breeding programs. To validate our findings, we conducted expression profiling of these paralogs and through computational analysis, elucidated the factors contributing to the diversity in the phenotypic traits observed in these rice accessions.

## Materials and Methods

### Plant material and growth conditions

Diverse rice accessions (272 numbers) belonging to the 3K Rice Genome panel were employed in this study (Supplementary Table S1). These accessions were phenotyped in response to N starvation condition at seedling stage for root related traits like primary root length (PRL), total root length (TRL), total root surface area (TRSA), total root volume (TRV), root average diameter (RAD), root tips and forks, for biomass such as shoot dry weight (SDW), root dry weight (RDW) and total biomass (TB) as well as N traits includes N concentration (N Conc.), total N uptake (TNUp), N use efficiency (NUE) and N acquisition efficiency (NAE) traits in the hydroponics system. The seeds were surface sterilized with 0.1% HgCl_2_ and rolled in germination paper and kept in the incubator at 28°C. After 5-6 days (emergence of coleoptile), the seedlings were transferred to nutrient solution with two N levels, low (0.01 mM) and sufficient (8.0 mM) using *PusaRicH* solution (Supplementary Fig. S1) (for composition of solution, refer to Sharma et al. 2018). The experimental set up was maintained at the National Phytotron Facility, IARI, New Delhi. The pH of *PusaRicH* solution was maintained between 4.9-5.1 using HCl or KOH. Seedlings were grown in two plastic containers (54 cm L x 41 cm W x 11 cm H) for each treatment, low and sufficient N, with four plants of each rice accession in one container. After 25 days, plants (n = 8) from each accession were sampled for biomass (shoot and root) and root traits. Roots were washed in distilled water. The length of longest root (primary root length) was measured manually using a ruler. Roots were scanned using a root scanner (Regent Instruments Inc., Canada) and the *.tif* images were analysed using WinRhizo Pro software. Dry weight of shoot and root tissues were recorded after drying samples in a hot air oven at 60℃ until a constant weight was reached. The tissue N% was analysed by Dumas method using a CHNS analyzer (Euro-Vector EA3000, Italy). The total N uptake was computed by multiplying N% with biomass per plant, while N acquisition efficiency (%) was calculated as the ratio of N content at low N to the N content at sufficient N. The physiological N use efficiency (mg dry weight mg N^-1^) was calculated as the ratio of total biomass to the total N uptake.

### Statistical analyses of phenotypic data

Analysis of variance (ANOVA) was carried out in AGRES software (Version 3.01). The principal component analysis (PCA), hierarchical cluster analysis, and correlation matrix were performed in the statistical software R version 4.2.2 (R Foundation for Statistical Computing, Vienna, Austria). A total of 14 traits were used for PCA; the components of Eigenvectors were identified for major traits contributing to the genetic variability for N uptake efficiency. These traits were considered for conducting hierarchical cluster analysis based on Ward’s methods which identified the contrasting accession groups. Graphs were plotted using GraphPad Prism version 5.01 (GraphPad Software, La Jolla, CA, USA).

### Haplotype analysis for *NRT1.1*

Haplotype analysis for three paralogs of *OsNRT1.1* gene was carried out by employing the in-built tool of SNP seek database. The analysis was used with default parameters and the Calinski criteria to determine the number of groups (k). The reference genome used for this study was Nipponbare. The haplotype analysis involved 272 rice accessions belonging to 12 subpopulations namely *ind2, aus, admix, indx, japx, aro, subtrop, ind1B, ind1A, ind3, temp*, and *trop*. Throughout the analysis, we employed the ’3kfiltered’ SNP set from the SNP Seek database, which had been curated using specific filtering criteria: (1) a minimum alternative allele frequency of 0.01, and (2) a maximum proportion of missing calls per SNP of 0.292 (http://snp-seek.irri.org/_download.zul). We utilized the pre-existing ’3kfiltered’ SNP set from the SNP Seek database and considered only non-synonymous SNPs and indels which alter the amino acid composition. The information on accession distribution among haplotypes and haplotype frequency was retrieved from the database. The phylogenetic tree based on SNPs was constructed in the SNP seek database and edited in MEGA 11. The gene structure of *OsNRT1.1* paralogs was prepared using http://wormweb.org/.

### Haplo-pheno association

The clusters obtained based on the phenotypic data were associated with haplotypes to determine the distribution of clusters among haplotypes. The pipeline used for the association of haplotype and phenotype data is summarised in Fig. 1. The efficient accessions belonging to each haplogroup were compared for all three genes separately to identify the superior haplotypes. The group with a higher mean trait value was reported as the superior haplogroup. A similar approach was also used to identify the inferior haplogroup. Further, expression analysis of *OsNRT1.1* paralogs were carried out in the selected rice accessions belonging to the superior and inferior haplogroup and were common in all three paralogs, that is, *OsNRT1.1a, OsNRT1.1b*, and *OsNRT1.1c*.

**Fig 1.**
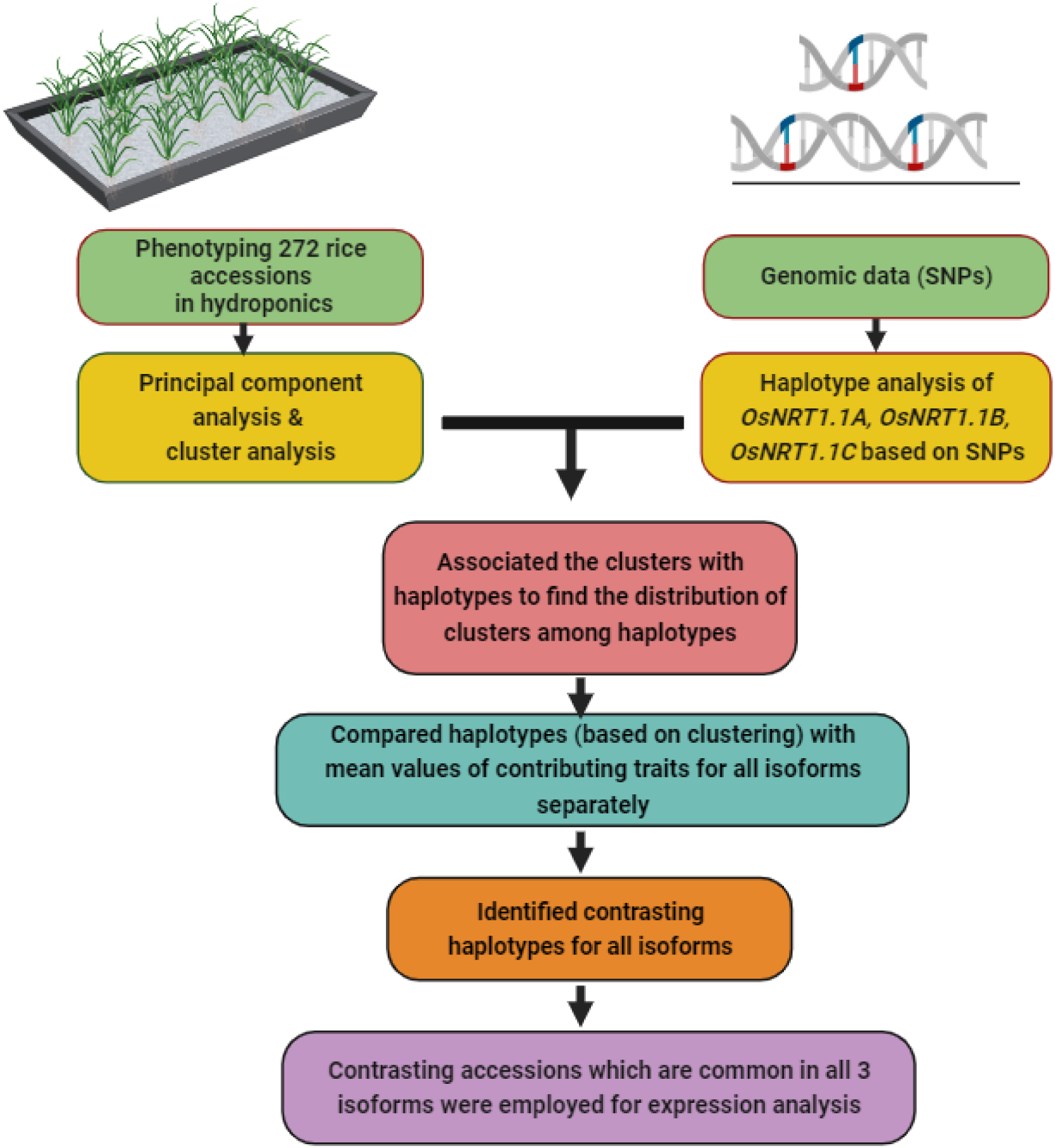
The pipeline depicting the association of haplotype and phenotype to find contrasting haplogroups of *OsNRT1.1* paralogs for further investigation.

### Expression analysis

For expression analysis of *OsNRT1.1* paralogs, 25-day-old plants of contrasting accessions were grown and exposed to low N and sufficient N conditions as described in the ‘Plant material and growth condition’. The shoot and roots were separated, immediately placed in liquid nitrogen, and kept at -80°C until the RNA was extracted. The RNA was isolated using the Invitrogen kit by Thermo Fisher Scientific followed by DNase treatment (Invitrogen kit) to eliminate genomic DNA contamination. The cDNA was synthesized using the High-Capacity cDNA Reverse Transcription kit (Thermo Fisher Scientific). The RT-qPCR was carried out using Brilliant III Ultra-Fast SYBR Green qPCR kit (Agilent Technologies) on an AriaMx G8830A Real-Time qPCR System (Agilent Technologies). The comparative cycle threshold method was used to calculate relative transcript levels at experimental conditions (low N) with three reference genes: *UBC 9, UBC 18* and *TIP41* (Sharma et al. 2021). Gene-specific primers were designed for *OsNRT1.1a, OsNRT1.1b*, and *OsNRT1.1c* using VectorNTI v11 software (Thermo Fisher Scientific). The set of primers was intentionally designed with or without SNPs found in either the forward or reverse primer sequences. Additionally, the length of the resulting amplicons was maintained within the range of 60 to 100 base pairs. The specificity of the primer was verified using the Primer-Blast website (https://www.ncbi.nlm.nih.gov/tools/primer-blast/index.cgi?GROUP_TARGET=on). The primers used for RT-qPCR and high-resolution melt curve (HRM) analysis are listed in Supplementary Table S2. The heat map for expression analysis was generated using TB tools (a <u>T</u>oolkit for <u>B</u>iologists integrating various biological data-handling <u>tools</u>)

### High-resolution melting curve analysis

The HRM analysis was carried out using MeltDoctor^TM^ HRM Master Mix on AriaMx G8830A Real-Time qPCR System (Agilent Technologies). Each 20 µl reaction comprised of 10 µl Evagreen HRM master mix, 7 µl distilled water, 1 µl each of forward and reverse primers and 1 µl cDNA (template). The reaction was carried out under the following conditions: 95℃ for 3 minutes (pre-denaturation), 40 cycles at 95℃ for 10 seconds, 60℃ for 12 seconds (amplification), and 90℃ for 2 minutes followed by 79℃ to 83℃ with a rise of 0.2℃ at each step (melt curve).

### Protein prediction and phylogenetic analysis

The protein sequences were aligned using MEGA 11. The protein blast analysis of OsNRT1.1 was conducted on the NCBI blast website (https://blast.ncbi.nlm.nih.gov/Blast.cgi). Homology modelling of proteins were employed using MODELLER 10.4 to determine the three-dimensional structure based on the known template protein structure. The quality of protein models was assessed using Procheck to generate a Ramachandran plot, as facilitated by the SAVESv6.0-Structure Validation Server (https://saves.mbi.ucla.edu/). Protein domains were predicted using the Conserved Domain Database (CDD) and represented using TB tools (https://www.ncbi.nlm.nih.gov/Structure/cdd/cdd.shtml). Protein motifs were predicted using the MEME Suite website (https://meme-suite.org/meme/). The predicted proteins were visualized in Pymol 2.5 software (https://pymol.org/2/). Finally, the active site of the proteins was predicted using CASTp 3.0 (http://sts.bioe.uic.edu/castp/index.html?1ycs).

## Results

### Identification of structural variations and nucleotide polymorphisms in *OsNRT1.1* paralogs

The full gene sequences of all three paralogs of *OsNRT1.1*, i.e., *OsNRT1.1A*, *OsNRT1.1B*, and *OsNRT1.1C*, were retrieved from the Rice Genome Annotation Project (RGAP) website (http://rice.uga.edu/) and a preliminary comparison was made. The schematic representation of *OsNRT1.1* on various chromosomes shows that *OsNRT1.1A*, *OsNRT1.1B*, and *OsNRT1.1C* are located on chromosomes 8, 10, and 3 respectively (Fig. 2A). The difference was found in the gene region and coding part of *OsNRT1.1* in all three paralogs. The CDS of *OsNRT1.1A, OsNRT1.1B,* and *OsNRT1.1C* was 1812 bp, 1791 bp and 1770 bp respectively. This is due to the conversion of a part of the coding DNA sequence (exon) into a non-coding sequence (intron) which is reflected in their protein sequences too (Fig. 2B). The 272 rice accessions were investigated for nucleotide variation in the sequences of *OsNRT1.1A*, *OsNRT1.1B*, and *OsNRT1.1C*. The nucleotide analysis of these paralogs revealed the presence of 2 non-synonymous (ns) SNPs and 420 indels in *OsNRT1.1A*, 2 nsSNPs and 213 indels in *OsNRT1.1B* whereas 5 nsSNPs and 102 indels in *OsNRT1.1C* with reference to Nipponbare. Most of the indels were identified in non-coding regions in all three paralogs which also could possibly alter the phenotype.

**Fig 2.**
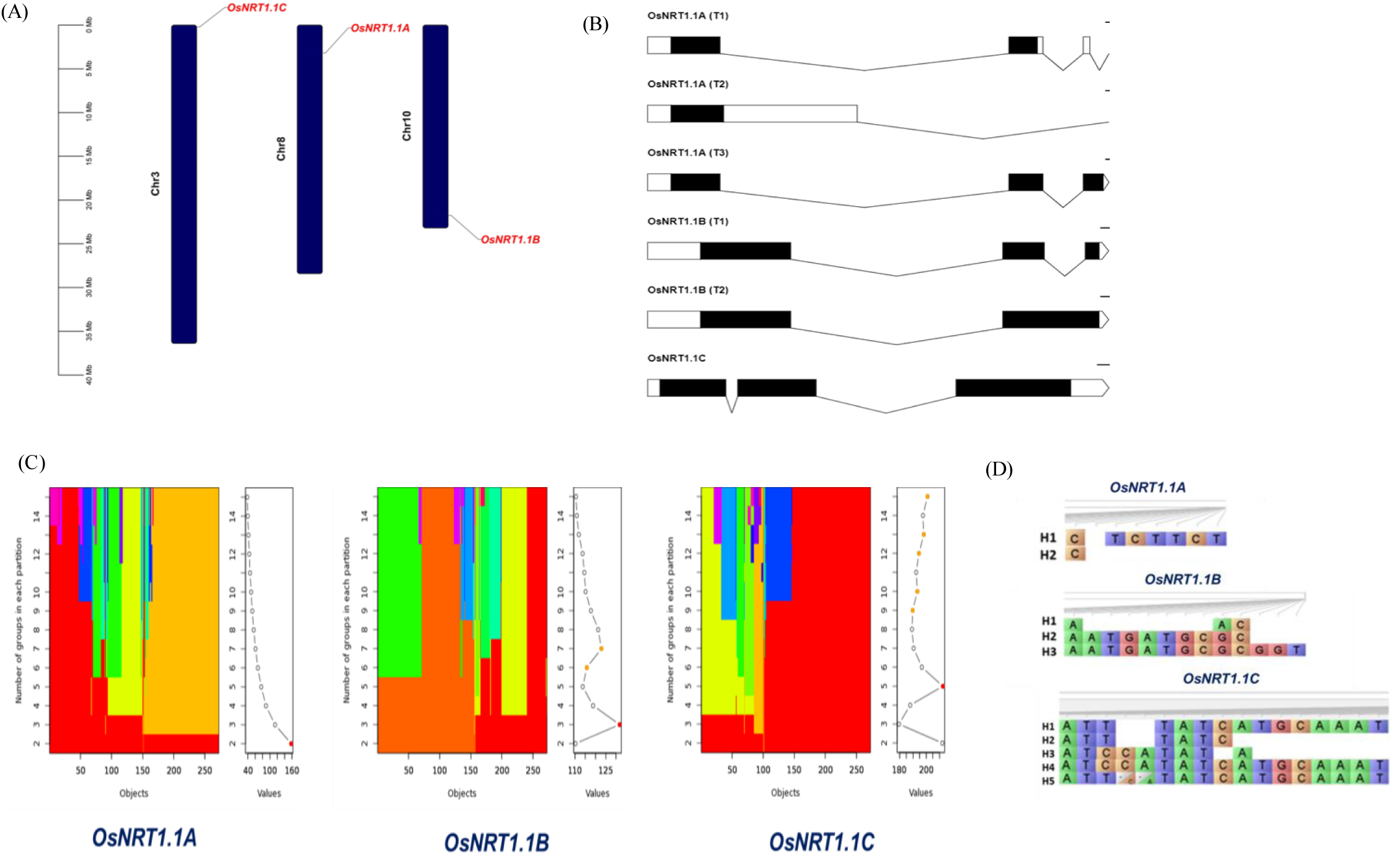
(A) Pictorial representation of localization of *OsNRT1.1* paralogs in rice chromosomes indicating *OsNRT1.1A* in chromosome 8, *OsNRT1.1B* in chromosome 10, and *OsNRT1.1C* in chromosome 3. (B) Summary of the total variations (gene length, CDS length, and protein length) detected in the *NRT1.1* paralogs. (C) Haplotype analysis of OsNRT1.1 paralogs across 272 rice accessions from the 3KRG panel. (D) Illustration of utilizing the Calinski criteria to ascertain the optimal K value for categorizing 272 rice accessions into haplotypes.

### Haplotype analysis of *OsNRT1.1*

The SNP data for 272 rice accessions were used for haplotype analysis for *OsNRT1.1* paralogs with Calinski criteria for k-group determination (Fig. 2C). According to Calinski criteria for *OsNRT1.1A*, the total 272 rice accessions were grouped into two haplotypes with 176 belonging to H1 and 96 accessions to H2 Fig. 2D. Similarly, for *OsNRT1.1B*, three haplotypes namely H1, H2 and H3 were identified each with 16, 156, and 100 accessions respectively. Five haplotypes (H1, H2, H3, H4 and H5) were observed in *OsNRT1.1C* each with 169, 20, 18, 52, and 13 accessions respectively. The accessions belonging to respective haplotypes are listed in Supplementary Table S3. Among the three paralogs, *OsNRT1.1C* displayed the highest genetic variability with five haplogroups, whereas *OsNRT1.1A* exhibited the lowest variability, with only two haplogroups. The H1 and H2 of *OsNRT1.1A* had 0.65% and 0.35% haplotype frequencies respectively. In case of the *OsNRT1.1B*, H1 had the least frequency (0.06%), followed by H3 (0.37%), and H2 (0.57%). In *OsNRT1.1C*, H1 had the highest frequency (0.62%), followed by H4 (0.19%), while other haplotypes possessed least frequencies (H2, H3, and H5 with 0.07%, 0.07%, and 0.05%, respectively).

### Phylogenetic study of *OsNRT1.1* paralogs

Phylogenetic analysis of the *OsNRT1.1* paralogs in rice revealed significant genetic divergence among various rice accessions. The distinctive haplotypes found in *O. sativa* accessions for all 3 paralogs of *OsNRT1.1* are represented by different colours (Fig. 3A-C). The phylogenetic tree revealed that certain accessions within one haplotype were positioned between accessions of another haplotype. This suggests a potential overlap or shared ancestry between these haplotypes. The overlapping possibly resulted from sequence similarity attributable to synonymous SNPs since as we utilized only indels and non-synonymous SNPs for haplotype analysis.

**Fig 3.**
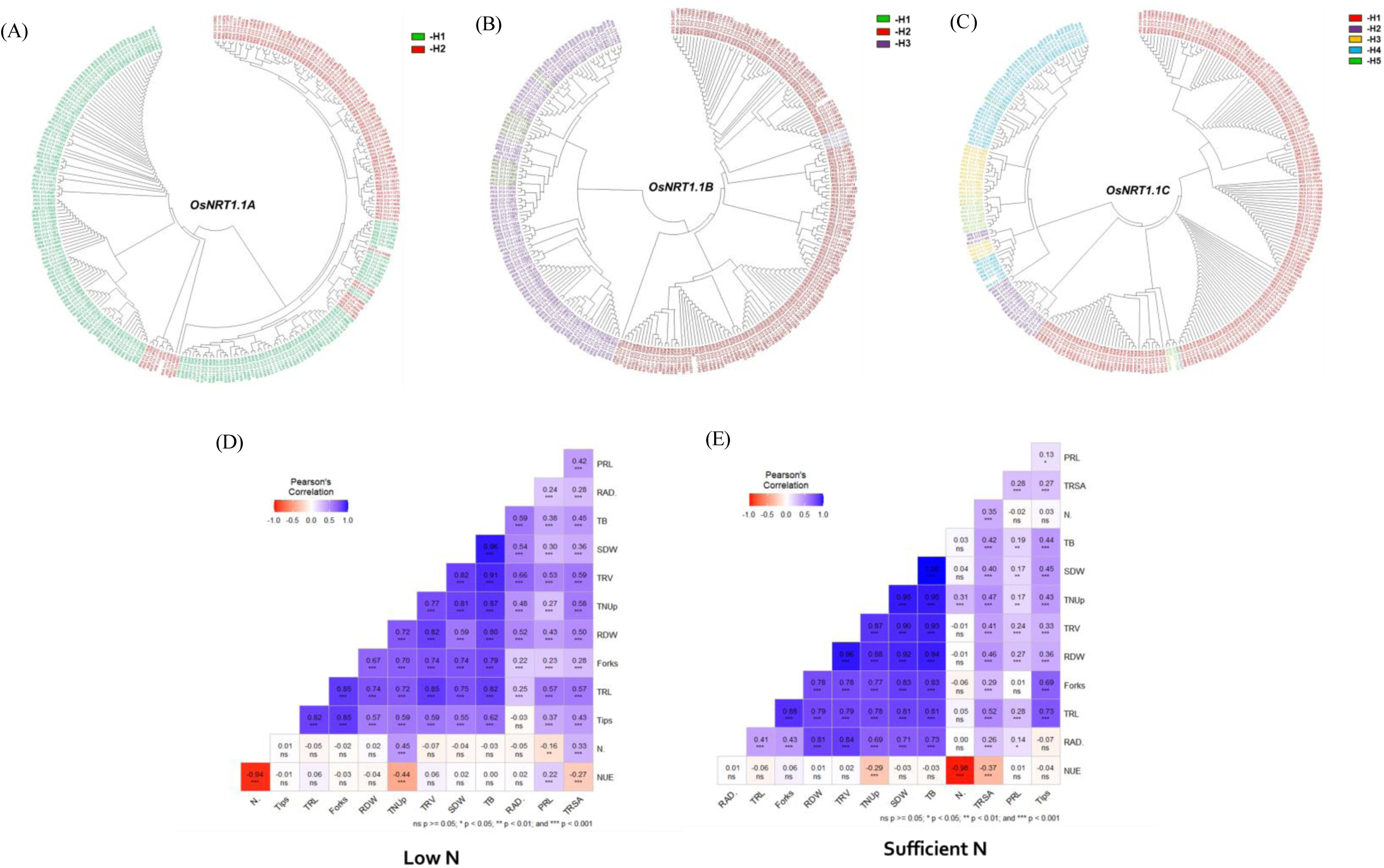
The phylogenetic tree was generated using the IRRI’s SNP search database’s built-in tool, and the haplotypes for (A) *OsNRT1.1A*, (B) *OsNRT1.1B*, and (C) *OsNRT1.1C* were manually coloured to represent them in distinct clusters. Linear correlation between various traits of 272 rice accessions in response to low N (D) and sufficient N (E) in hydroponics. Abbreviation: PRL-primary root length, SDW-shoot dry weight, RDW-root dry weight, TB-total biomass, TRL-total root length, TRSA-total root surface area, TRV-total root volume, RAD-root average diameter, root tips, root forks N. Conc-N concentration (N%), TNUp-total N uptake, and NUE-nitrogen use efficiency.

### Phenotyping rice accessions for nitrogen uptake

The *NRT1.1* is a key nitrate transporter associated with environmental stress, particularly under low N conditions. The root traits varied significantly (P<0.01) among the rice accessions under different N treatments (Supplementary Table S4). In comparison to control, the low N supply resulted in significant reduction in SDW (43.5%), RDW (5.4%), and TB (37.1%). Similar results were observed for root traits under low N conditions, PRL increased by 20.0%, whereas significant reduction was observed in TRSA (5.9%), TRL (10.2%), TRV (17.8%), root tips (32%) and forks (35%) as compared to control. However, RAD showed no significant differences between the treatments. The N related traits like N concentration (46.3%) and TNUp (66.4%) was significantly reduced, but NUE increased by 94.5% under low N condition (Supplementary Fig. S2 A to N). The Pearson’s correlation coefficient revealed that TB, SDW, and RDW were significantly positively correlated with root traits including TRSA, TRV, RAD, and TRL under low N and sufficient N conditions (Fig. 3 D, E). Similarly, TNUp was significantly positively correlated with all root and biomass traits, but was negatively correlated with NUE. However, PRL was associated positively with NUE and negatively with N% at low N stress.

### Principal component analysis and cluster analysis

The principal component analysis (PCA) revealed that traits like RDW, SDW, TB, TRSA, TRL, TRV, N Conc., TNUp, NUE contributed to the genotypic variability (Fig. 4A). Further, hierarchical clustering based on Ward’s method was carried out with 9 contributing traits resulting in three clusters. By comparing the mean values of 9 traits between the clusters, efficient (39 accessions), inefficient (125) and intermediate (108) clusters were identified (Fig. 4B and Supplementary Table S5). The inefficient cluster exhibited a marked reduction (61-70%) in SDW, RDW, TB, TRL, TRSA, and TRV compared to the efficient cluster. Conversely, TNUp and NUE were also reduced by 57% and 17% respectively, while N Conc. increased by 14% compared to the efficient cluster (Supplementary Fig. S3).

**Fig 4.**
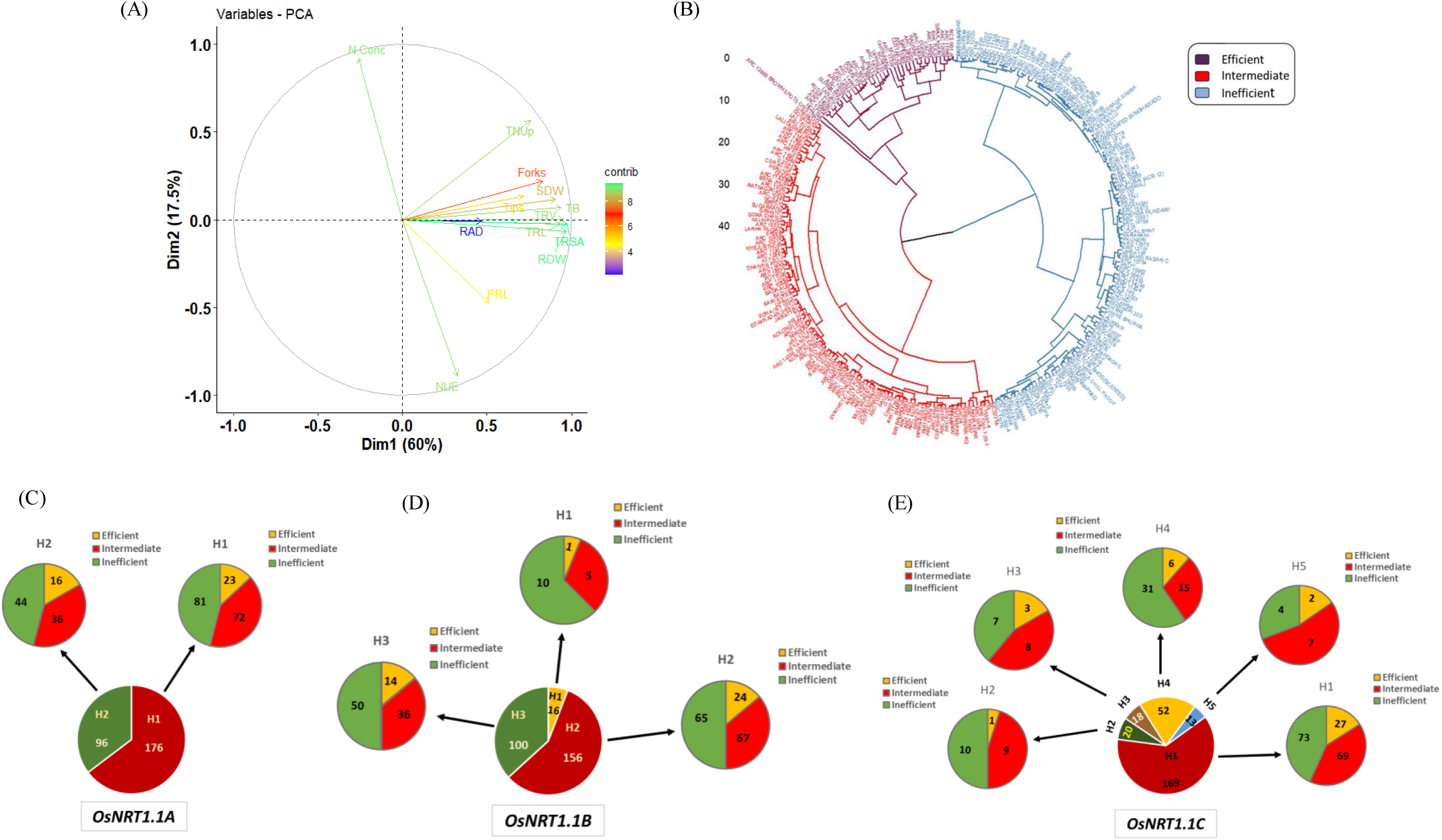
(A) Principal component analysis to identify the most contributing traits governing genotypic variability in response to N stress. Green indicates the most contributing traits and purple indicates the least contributing traits. (B) Hierarchical clustering analysis of 272 rice accessions performed with nine contributing traits resulted from PCA. Three clusters represented as N-efficient, N-intermediate, and N-inefficient. Association of (C) *OsNRT1.1A*, (D) *OsNRT1.1B*, and (E) *OsNRT1.1C* haplotypes with different clusters obtained from the phenotype data.

### Haplo-pheno association

The results from hierarchical cluster analysis using phenotypic traits were associated with haplotypes based on SNP analysis of *OsNRT1.1* in order to find the distribution of clusters among haplotypes. For *OsNRT1.1A,* out of 39 N-efficient accessions, H1 and H2 possessed 23 and 16 accessions respectively (Fig. 4C). Among 108 intermediate accessions, H1 had 72 and H2 had 36 accessions whereas in the N-inefficient cluster (125), 81 belonged to the H1 haplotype and 44 fall into H2 haplotype. Similarly, for *OsNRT1.1B*, H1 possessed only one accession present in N-efficient cluster, five in N-intermediate and 10 in the N-inefficient clusters (Fig. 4D). In H2, 24 accessions were present in the N-efficient cluster, 67 in N-intermediate, and 65 in the N-inefficient clusters. Further, the H3 haplotype possessed 14 accessions in the N-efficient, 36 in N-intermediate, and 50 in the N-inefficient clusters. However, in *OsNRT1.1C,* out of 169 accessions in H1, 27 were N-efficient, 69 were N-intermediate and 73 were N-inefficient (Fig. 4E). Similarly, in H2 with 20 accessions, one was N-efficient, nine N-intermediate, and ten were N-inefficient accessions. H3 contained 18 accessions, with 3, 8 and 7 belonging to N-efficient, N-intermediate and N-inefficient respectively. The haplotype H4 comprised of 52 accessions, with 6, 15 and 31 belonging to N-efficient, N-intermediate, and N-inefficient clusters. Lastly, H5 possessed 13 accessions, with two N-efficient, seven N-intermediate, and four N-inefficient accessions. This information clearly depicts the distribution of accessions based on their N-efficiency levels within haplogroups of the *OsNRT1.1* paralogs.

### Identification of superior and inferior haplotypes

To identify the superior haplotypes, the mean values of accessions belonging to the N-efficient cluster under haplotypes H1 and H2 were compared with their performance under low N conditions. Similarly, to identify the inferior haplotype, the mean values of accessions belonging to N-inefficient cluster under the H1 and H2 were compared with the least performance under low N (Supplementary Fig. S4 A to G). For example, in *OsNRT1.1A*, the mean of accessions for shoot dry weight under low N in H1 (111.5 mg plant^-1^) and H2 (127.7 mg plant^-1^) under the N-efficient cluster were compared for their performance which clearly showed that the haplotype H2 was superior (Supplementary Table S6). Likewise, in the N-inefficient cluster, the mean value of shoot dry weight at low N for H1 (78.7 mg plant^-1^) was lesser than H2 (80.0 mg plant^-1^), thus, the former was marked as an inferior haplotype.

In a similar manner, the superior and inferior haplogroups were identified depending on the most contributing traits for *OsNRT1.1B* (Supplementary Table S7) and *OsNRT1.1C* (Supplementary Table S8). In *OsNRT1.1C*, limited accession availability in some haplotypes led to the selection of two superior and two inferior haplotypes. In conclusion, it has been identified that the superior haplotypes (SH) were H2 in *OsNRT1.1A*, H3 in *OsNRT1.1B*, and H3, H1 in *OsNRT1.1C* among the N-efficient cluster. The inferior haplotypes (IH) were H1 in *OsNRT1.1A*, H3 in *OsNRT1.1B*, and H3, H2 in *OsNRT1.1C* among the N-inefficient cluster. In *OsNRT1.1A*, H1 (IH) demonstrated a significant reduction of 67-74% in SDW, RDW, and TB (Fig. 5A), as well as 64-77% reductions in TRL, TRSA (Fig. 5B), and TRV. However, TNUp and NUE exhibited 60.2% and 20.7% reductions, whereas N Conc. increased by 22.2%, compared to H2 (SH) relative trait values. Similarly, in *OsNRT1.1B*, compared to H3 (SH) relative trait values, H3 (IH) exhibited reductions of 63 to 72.5% in SDW, RDW, and TB (Fig. 5C), along with 66 to 75% reductions in TRL, TRSA (Fig. 5D), and TRV. Additionally, TNUp and NUE showed reductions of 59% and 14.2%, while N Conc. increased by 11.8%. In case of *OsNRT1.1C*, H3 (IH) showed significant reductions in SDW, RDW, and TB by 75-82% (Fig. 5E), and in TRL, TRSA, and TRV by 75-86% (Fig. 5F). Also, it exhibited a reduction in TNUp (72%) and NUE (20.1%), with a notable increase in N Conc. (21.6%) compared to H3 (SH). Likewise, H2 (IF) displayed reductions in SDW, RDW, and TB (66-73%) as well as TRL, TRSA, and TRV (58-73%), alongside a reduction in TNUp (59.4%) and NUE (23.2%), coupled with a 24% increase in N Conc. compared to H1 (SH) relative trait values.

**Fig 5.**
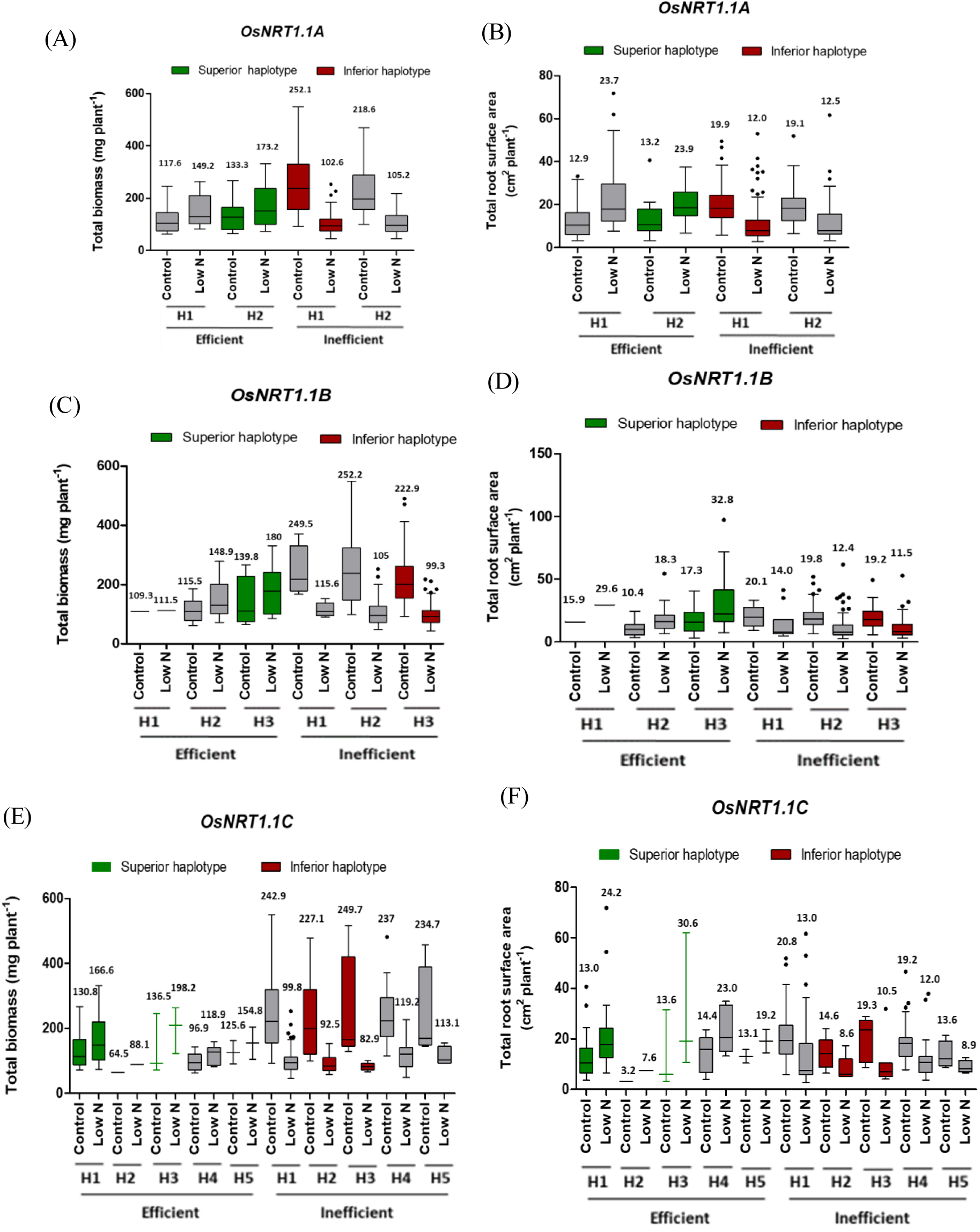
Association of haplotypes of *OsNRT1.1A*, *OsNRT1.1B*, and *OsNRT1.1C* with total biomass (A, C, E) and total root surface area (B, D, F) of 25 days old rice seedlings grown with low and sufficient N supply. Outliers are mentioned as dots. The value above the bar is the mean trait value. Red highlights the inferior haplotype while green highlights the superior haplotype.

### Identification of accessions carrying superior and inferior haplotypes

Accessions with superior haplotypes of all three *OsNRT1.1* paralogs as a result of haplo-pheno association could be valuable for haplotype-based breeding. A total of 5 accessions were found to have the superior haplotypes of all three paralogs of *OsNRT1.1* namely ARC 7091, ADT 12, ARC 11571, SIMUL KHURI and ARC 12920. However, among the inferior haplotypes, three accessions were found to be common in all, such as KARAHANI, ARC 13591 and BK 26 (Fig. 6A). These contrasting accessions were further utilized for validation. **Validation by expression analysis**

**Fig 6.**
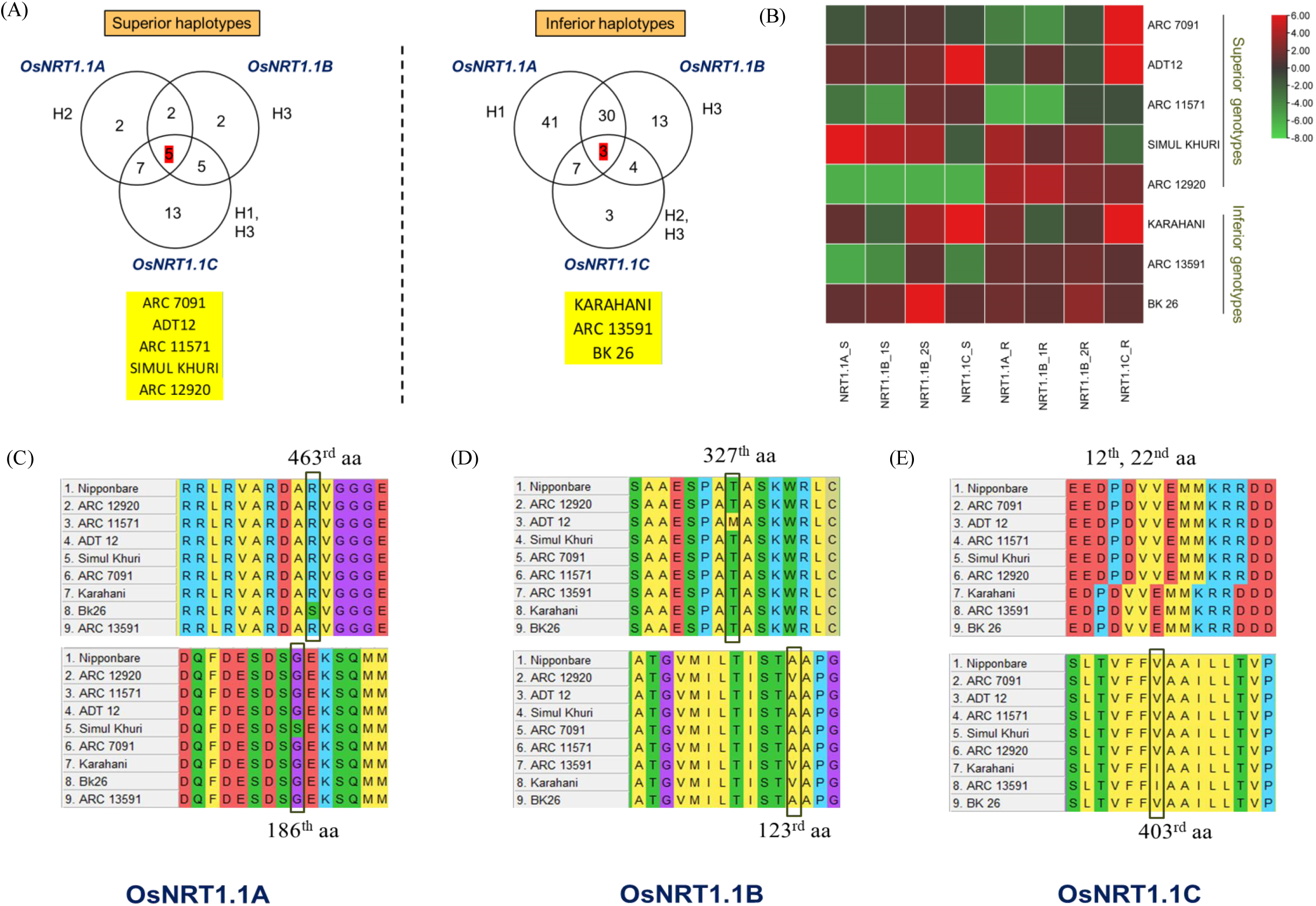
(A) Selection of accessions that are common in superior and inferior haplogroups of *OsNRT1.1* paralogs as represented by Venn diagram. (B) Heatmap depicting the relative expression of *OsNRT1.1* paralogs in superior and inferior haplogroups. Multiple sequence alignment of (C) OsNRT1.1A, (D) OsNRT1.1B, and (E) OsNRT1.1C proteins shows amino acid changes between the contrasting rice accessions.

The selected contrasting accessions were investigated for expression analysis of *OsNRT1.1* paralogs. Among the accessions, SIMUL KHURI exhibited the highest expression in both shoots and root tissues for *OsNRT1.1A* and *OsNRT1.1B* but, *OsNRT1.1C* was downregulated. Out of five accessions in superior haplogroups, three showed no-significant difference in the expression of OsNRT1.1. Interestingly, the relative expression of OsNRT1.1C increased by 16 and 18-fold in ARC 7091 and ADT 12, respectively (Fig. 6B). The superiority of these accessions could be linked to OsNRT1.1-mediated nitrate signalling. In a few accessions belonging to inferior haplogroup, it was noted that the gene expression was usually higher in the root but not in the shoot, which might account for its inefficiency. Interestingly, ARC 12920 (superior genotype) had a similar pattern of expression with ARC 13591 (inferior genotype) in both shoots and roots (Supplementary Fig. S5 A to H). Further investigation of ARC 12920 reveals that it possesses lower values of various traits under sufficient N but higher values under low N conditions, placing it in the N-efficient cluster. However, no clear pattern defining the superior and inferior accessions could be seen. The gene expression results suggest the equally important role of post-translational regulation and amino acid change in a protein.

### Comparative protein sequence analysis in contrasting accessions

The changes due to SNPs in the coding sequences (CDS), the amino acid sequences of OsNRT1.1 were compared to determine the rationale for the accession’s efficiency and inefficiency to N. Notably, in OsNRT1.1A protein, SIMUL KHURI and BK 26 displayed amino acid changes at positions 186 (serine instead of glycine) and 463 (serine instead of arginine), respectively (Fig. 6C). Within OsNRT1.1B, KARAHANI, ARC 13591, and ARC 12920 exhibited substitution of threonine residue with cysteine at position 123 (Fig. 6D), situated in a conserved domain and motif region of the protein (Supplementary Fig. S6 A, B). Conversely, ADT 12 had an alteration at position 327 (threonine substituted by methionine), unique to the conserved domain. In OsNRT1.1C, inferior accessions KARAHANI and ARC 13591 showed an amino acid change at position 403 (isoleucine replacing valine), present in both the conserved domain and motif regions (Fig. 6E and Supplementary Fig. S6 A, B).

### Unveiling Protein structure-function relationships

Protein structures corresponding to OsNRT1.1A, OsNRT1.1B, and OsNRT1.1C were examined for structural variations caused by amino acid alterations. According to the Ramachandran plot, more than 99.2% to 99.6% of the residues in the OsNRT1.1 proteins were found in the core and allowed region, whereas just two to four residues were found in the disallowed region. The position of amino acid change in the transmembrane helix (TMH) is represented in the 3D-protein structure. In OsNRT1.1A, the 186^th^ aa change was found in the TMH5 loop and the 463^rd^ aa change was present in TMH 9-10 loop (Fig. 7A). Simultaneously, the 123^rd^ and 327^th^ aa changes in OsNRT1.1B were found in the TMH3 and TMH6-7 loop of the protein, respectively (Fig. 7B). However, in OsNRT1.1C, the 403^rd^ aa change was present in the TMH 8, whereas the 12^th^ aa deletion (glutamic acid, E) and 22^nd^ aa insertion (aspartic acid, D) were present in the N terminal region of the protein (Fig. 7C). However, among all the amino acid changes in OsNRT1.1, isoleucine was substituted at 403^rd^ position in OsNRT1.1C that is present in the active site of the protein, which was specific to the identified inferior rice accessions (Fig. 7D). This amino acid modification noticeably affects the active site (AS) in inferior accessions (AS volume: 4473.4 Å3) as compared to the superior accessions (AS volume: 7423.1 Å3), which might have accounted for the N-inefficiency of these accessions.

**Fig 7.**
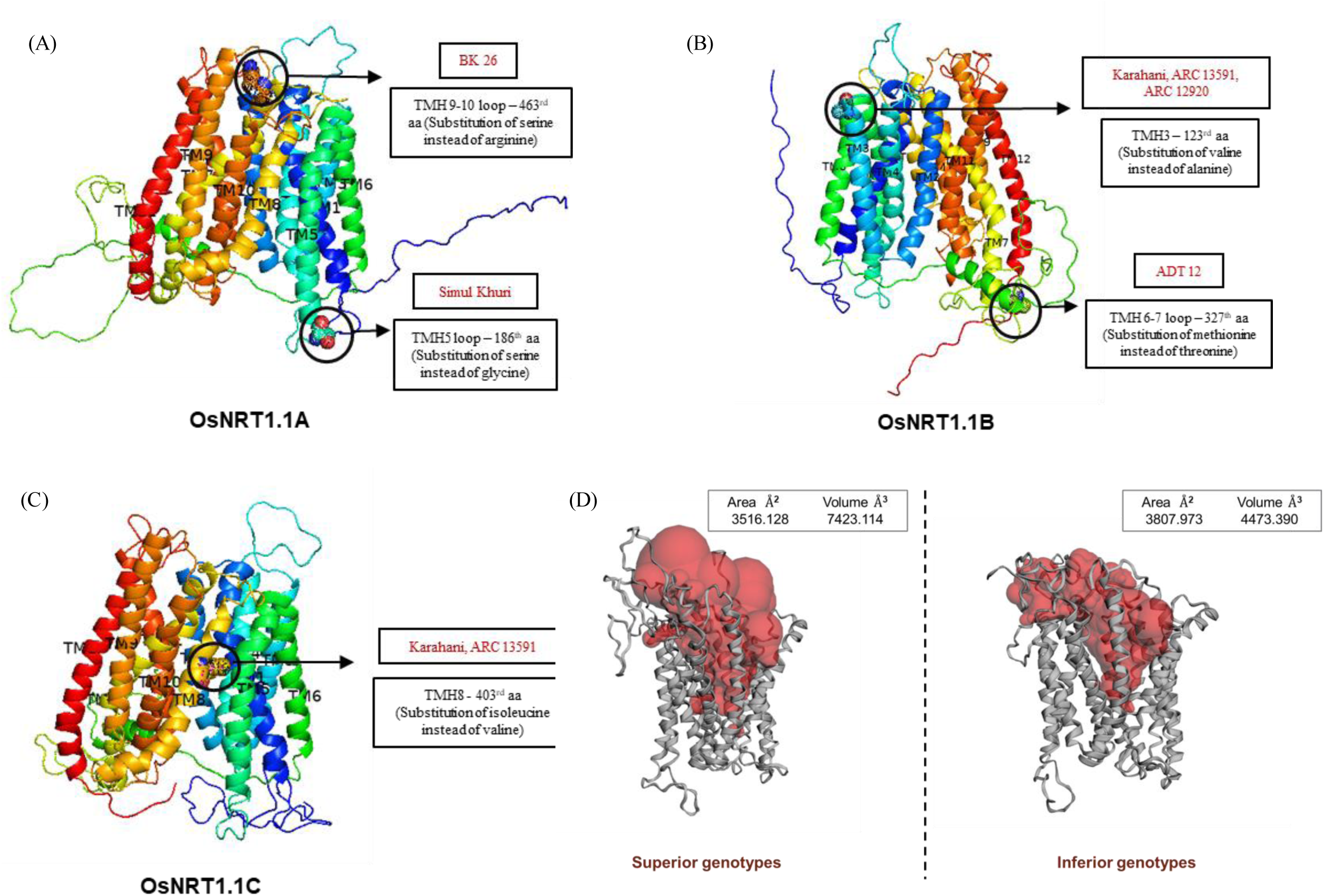
Homology-modelled proteins of (A) OsNRT1.1A, (B) OsNRT1.1B, and (C) OsNRT1.1C represented with the localization of amino acid changes in the protein. (D) Schematic representation of active site changes in OsNRT1.1C protein in superior and inferior genotypes due to SNP changes.

### Validation of mutation in OsNRT1.1C by HRM analysis

Variations among the 3 different accessions in OsNRT1.1C were identified by analysing temperature-shifted curves. The melting curve differences relative to the selected accession as baseline or accession without mutation is illustrated in Fig 8. The superior accession, SIMUL KHURI with GG homozygous SNP is taken as a baseline. However, the inferior accession ARC 13591 with heterozygous SNP (G/A) and KARAHANI with homozygous SNP (AA) accession depicts the polymorphism in these accessions through the shapes of the melting curves, highlighting the visual differentiation among all accessions.

**Fig 8.**
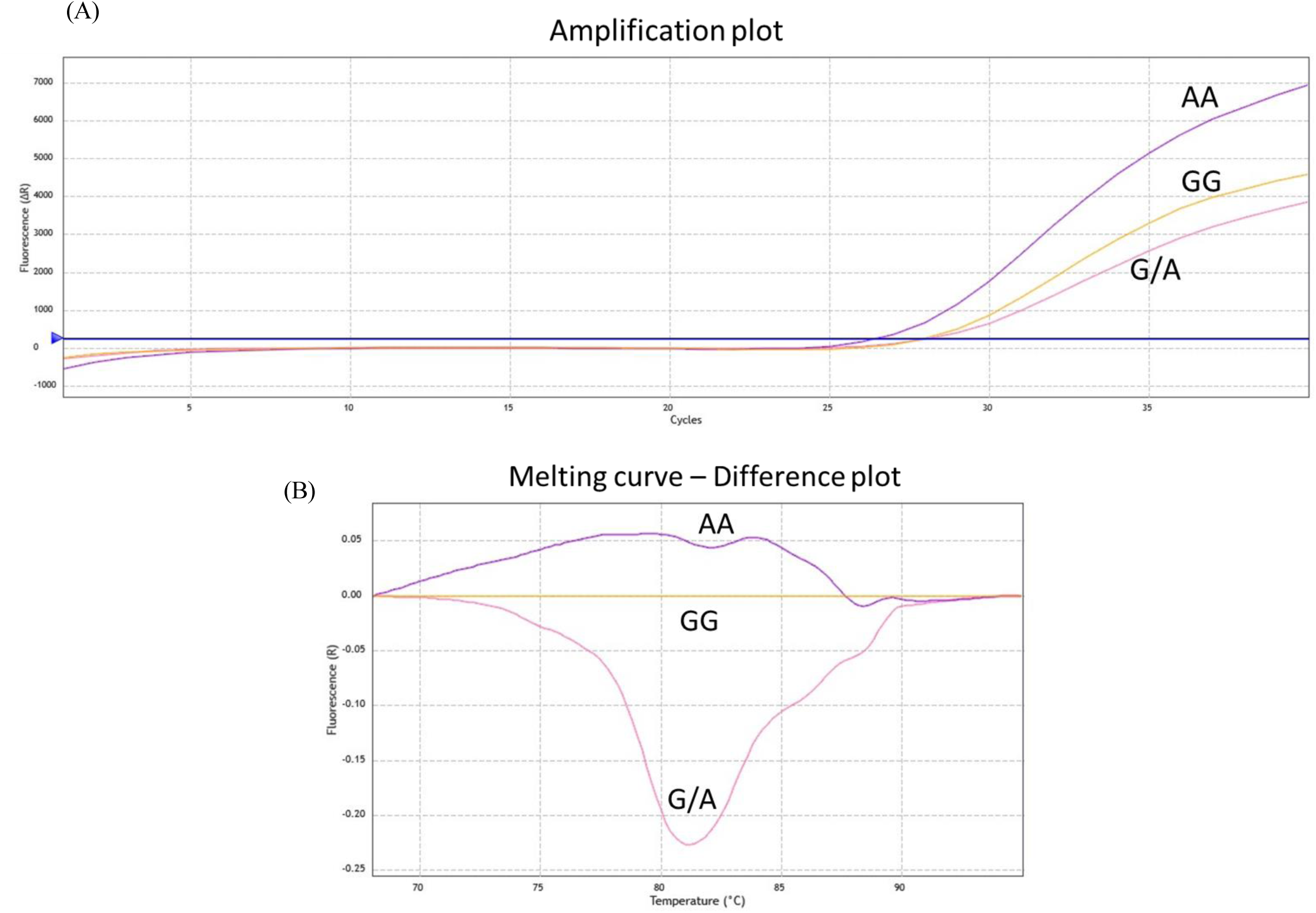
Discrimination of superior and inferior accessions based on an SNP in OsNRT1.1C. Amplification plot (A) and melting curve difference plot (B) depict the melt curve variation between the accessions based on SNPs.

## Discussion

In this study, we used 272 rice accessions from the 3KRG panel, a rich repository of genetic diversity, to identify the significant SNPs which account for their superiority at low N condition conferred by the potential candidate gene, *NRT1.1* (Ye et al. 2019). Arabidopsis *AtNRT1.1* (*CHL1/AtNPF6.3*) besides associated with nitrate uptake and transport as well as nitrate-sensing “transceptor’’ (Sun et al. 2014), it also exhibits auxin transport capabilities and influences nitrate-dependent root development, suggesting multi-functionality (Wang et al. 2020). Thus, finding the genotype with a superior haplotype of *OsNRT1.1* would be valuable for haplotype-based breeding. Similar report on haplotype analysis of six QTLs in rice that were strongly associated with early root vigour, was used to identify the donors with superior haplotypes (Anandan et al. 2022). *SERINE CARBOXYPEPTIDASE I* (*Cxp1*), a potential gene for malting quality, is regulated in *cis*, and the presence of various SNP haplotypes within the gene was correlated with the expression level of the gene (Potokina et al. 2006). In our study, among the rice accessions, the ones exhibiting maximum root growth, biomass, and N-related traits under N deficiency offers a promising opportunity to introduce this novel allele into the elite cultivars. This genetic transfer holds the potential for enhancing N starvation tolerance in rice varieties.

Population genetics and evolutionary factors, such as mutation and recombination rates, and selection, all affect the haplotypic variation of a specific region (Zaitlen et al. 2005). In this context, our findings highlight a notably low haplotype diversity within *OsNRT1.1A* and *OsNRT1.1B* across the 272 accessions predominantly originating from India. This suggests a limited genetic diversity within this population. Exploring rice populations from different geographic regions, characterized by a broader genetic diversity, is anticipated to yield a wider array of haplotypes, offering valuable resources for the discovery of superior haplotypes on a global scale. Additionally, haplotypes with a lower frequency in rice for *NRT1.1* genes are relatively rare variants within the gene pool, representing a smaller portion of the overall genetic diversity in rice populations. Exploring haplotypes of additional genes related to N uptake and utilization would provide a more holistic understanding and ultimately lead to the identification of a superior genotype with enhanced N-related traits.

### Nitrogen-driven variation in phenotypic pattern

Present study depicts significant variation in phenotype i.e., root morphological traits, biomass, and N-related traits with respect to different N levels. The observed outcome aligns with previous studies on various crops investigating the influence of N on root morphology, biomass, and N-related traits, reinforcing the consistency of these findings in existing research (Zhao et al. 2005; Kun et al. 2014; Zhu et al. 2022; Tiwari et al. 2023). Our results also showed a substantial increase in primary root length under low N stress, a key trait aiming to enhance nitrate acquisition. In contrast, root length decreased under sufficient N conditions but the total root surface area increased along with more number of lateral roots thereby increasing the root biomass under sufficient conditions for effective N uptake (Supplementary Fig. S3). When subjected to low N supply, inefficient accessions exhibited reduced total biomass, primarily attributed to their inefficient N uptake mechanisms. Reduction in NUE at higher N rates indicated that rice plants were either unable to absorb or utilize N at higher rates, or that N loss was greater than the rate of N uptake (Fageria et al. 2005). Our results also confirmed that at lower N rates, NUE increases and at higher N rates NUE decreases. This suggests that the absorption mechanisms in rice seedlings may have been saturated, making them unable to take N when it was given in excess.

### N-efficiency in rice accessions is governed by differential expression of *OsNRT1.1* alleles

For plants to survive better in low N environments, *NRT1.1* expression needs to be increased in shoots rather than roots. The increase in *NRT1.1* expression in shoots is polymorphism-dependent and affects a range of N deprivation responses in Arabidopsis (Sakuraba et al. 2021). In this context, the accessions ARC 13591 and KARAHANI belonging to inferior haplogroup, showed downregulation of *OsNRT1.1B* and all *OsNRT1.1* paralogs in shoots, which is a crucial factor for plant survival. This disparity in expression levels in these accessions might have led to the N-inefficiency (Fig. 6B). Compared to *AtNRT1.1* and *OsNRT1.1B*, which primarily participate in the initial nitrate-triggered response, *OsNRT1.1A* appears to have a more pivotal function in efficiently regulating N utilization, especially under conditions of relatively stable N supply encompassing both ammonium and nitrate sources (Wang et al. 2018). Further, it was shown that *OsNRT1.1A* was expressed in both roots and shoots when rice plants were grown in the presence of both nitrate (NO_3_^−^) and ammonium (NH_4_^+^). On the other hand, *OsNRT1.1B* exhibited higher expression levels mainly in the roots in the presence of nitrate (NO_3_^−^), while the expression of *OsNRT1.1C* was minimal in both roots and shoots (Vera et al. 2021). Our findings revealed intriguing variations in gene expression patterns in contrasting accessions. Notably, higher levels of *OsNRT1.1C* expression were noted in the roots of all the accessions except ARC 11571 and SIMUL KHURI and shoots of ADT 12, KARAHANI, and BK 26. Conversely, *OsNRT1.1B* exhibited higher expression in shoots of all the accessions except ARC 12920 and roots of SIMUL KHURI, ARC 12920, ARC 13591, KARAHANI, and BK 26. Additionally, our results highlighted substantial variations in the expression levels of the two transcripts (mRNA splicing products) of *OsNRT1.1B_1* and *OsNRT1.1B_2* across the selected accessions, highlighting the complexity of the regulatory mechanisms of these genes (Fig. 6B). These results emphasize the need for further research to elucidate the underlying factors contributing to the expression variations in different *OsNRT1.1* alleles.

### OsNRT1.1C, a potential player regulating nitrogen uptake in plants

Any change in amino acid sequence can impair protein function due to disruptive folding and protein structure. These changes may lead to loss of enzymatic activity, altered binding properties, or even protein degradation, ultimately, impacting vital biological processes in plants. We demonstrated that except an insertion at position 12^th^ and deletion at 22^nd^ in OsNRT1.1C, all alterations in amino acid sequences found in all three OsNRT1.1 variants were located within the conserved domain, and a few in the motifs, which potentially impacted the functionality (Supplementary Fig. S6). These changes in the conserved regions can disrupt essential functional elements, thus influencing the overall functionality of OsNRT1.1 protein. A Cu-sensitive Arabidopsis accession, Chisdra-2, has a unique CPC(x)6P domain which is a strictly conserved. The substitution of leucine with proline resulted in loss of function suggesting that the functional integrity of the Cu-translocating ATPase, HMA5 (HEAVY METAL TRANSPORTING ATPases5), regulates a portion of diversity in Cu tolerance in Arabidopsis (Kobayashi et al. 2008). However, SNPs and INDELs at non-coding regions could also potentially alter the phenotype of the plants. A SNP mutation (G to A) at the splice acceptor site of the intron in *GA3ox* resulted in altered splicing, leading to two isoforms in dwarf plants of watermelon (*Citrullus lanatus*) - one with intron sequences retained and the other with a 13-bp deletion in the second exon, both of which resulted in truncated proteins and loss of the functional Fe2OG dioxygenase domain (Sun et al. 2020). Hence, the exploration of INDELs present at non-coding regions of *OsNRT1.1* paralogs could potentially uncover the mysteries surrounding this gene.

Single amino acid change can also influence the functionality of proteins. For example, the distinct Pyrabactin selectivity exhibited by PYL1 and PYL2, is determined by a single amino acid alteration, switching between valine and isoleucine in the corresponding position demonstrated in Arabidopsis (Yuan et al. 2010). Another example demonstrated by replacing Val_566_ with Ile_566_ showed the potential to interfere with hydrogen bonding and alter the solvent accessibility of 22 amino acid residues, resulting in the formation of a brief α-helix at the C-terminus of Anti-Müllerian hormone (AMH) in chickens (Dang et al. 2020). While both valine and isoleucine are hydrophobic amino acids, isoleucine possesses a more substantial hydrophobic nature due to its larger side chain with additional methylene groups, resulting in a greater volume compared to valine (Behmard et al. 2012). Our findings are consistent with these reports, demonstrating that the substitution of isoleucine for valine at the 403^rd^ position has a pronounced effect on the active site of OsNRT1.1C protein. This alteration significantly disturbs the spatial arrangement and chemical properties of the active site, potentially affecting its binding affinity with the substrate (most probably nitrate) and protein function (Fig 7D). These observations emphasize the importance of specific amino acid residues in modulating the protein function, thus, shedding light on the functional intricacies of the OsNRT1.1C.

Both apo and nitrate-bound AtNRT1.1 crystal structure as well as *in vitro* binding and transport studies point to the significant role of His_356_ in nitrate (substrate) binding (Parker and Newstead 2014). Surprisingly, His_356_ is not well conserved in NRT1 members in rice other than OsNRT1.1B, indicating that additional NRT1 members transport and recognize nitrate using an unidentified mechanism. So, additional investigation into the molecular mechanisms underlying nitrate transport in OsNRT1.1A and OsNRT1.1C could provide insights into how changes in amino acids at the protein level contribute to the N-inefficiency observed in these accessions. The subcellular distribution of OsNRT1.1A within the vacuole (Wang et al. 2018) and the presence of OsNRT1.1B in the plasma membrane (Wang et al. 2020) are well-documented (Supplementary Fig. S7) but there is a lack of information regarding the localization of OsNRT1.1C. This gap in knowledge highlights the importance of further research on OsNRT1.1C which will provide valuable insights of its role in nitrate transport. Thus, our findings strongly suggest that investigating OsNRT1.1C could contribute significantly to the understanding of nitrate transport and utilization mechanisms in plants.

## Author contributions

D.E. haplotype analysis, phenotyping experiment, data analysis, initial manuscript draft; R.P. conceptualization, planning, interpretation, edited and finalised the manuscript, resources, supervision, fund acquisition; S.S. gene expression & HRM analysis, assisted in phenotyping data analysis; T.K. nitrogen analysis; A.D., B.B., R.K.E. assisted in haplotype data analysis; N.A. provided purified seed material; S.K. conceptualization, planning. All authors read and approved the manuscript.

## Supplemental data

The following materials are available in the online version of this article.

**Supplementary Figure S1**. Pictorial representation of rice plants grown in sufficient and low N at 2 DAT, 10 DAT, 18 DAT, and 25 DAT.

**Supplementary Figure S2**. Phenotypic characterization of 272 rice accessions for various traits (A) shoot dry weight (B) root dry weight (C) total biomass (D) primary root length (E) total root length (F) total root volume (G) total root surface area (H) root average diameter (I) root tips (J) root forks (K) N concentration (%) (L) Total N uptake (M) N use efficiency (N) N acquisition efficiency among the phenotype characterization of 272 rice genotypes.

**Supplementary Figure S3.** Comparisons of various clusters obtained by hierarchical cluster analysis of 272 rice accessions for various traits. (A) shoot dry weight (B) root dry weight (C) total biomass (D) total root length, (E) total root surface area (F) total root volume (G) total N uptake and (H) N use efficiency.

**Supplementary Figure S4.** Association of haplotypes with phenotypic clusters for various traits in 272 rice accessions grown under low N and sufficient N supply (A) shoot dry weight, (B) root dry weight, (C) total root length, (D) total root volume, (E) nitrogen concentration, (F) total nitrogen uptake, and (G) nitrogen use efficiency. Outliers are mentioned as dots. The value above the bar is the mean trait.

**Supplementary Figure S5.** Relative expression analysis of *OsNRT1.1* paralogs in contrasting accessions. Duncan analysis was employed to test the statistical significance of expression results.

**Supplementary Figure S6.** Schematic representation of the distribution of various (A) conserved domains and (B) motifs in OsNRT1.1 protein paralogs. The location of amino acid change in the motif is marked.

**Supplementary Figure S7.** Subcellular localization of *OsNRT1.1A* in the plasma membrane and *OsNRT1.1B* in the vacuole.

**Supplementary Table S1**. List of 272 rice accessions used for haplotype and phenotype analyses.

**Supplementary Table S2.** List of primers used for RT-qPCR to validate the transcript levels of *OsNRT1.1* paralogs.

**Supplementary Table S3**. Rice accessions belonging to each haplotype of *OsNRT1.1* paralogs.

**Supplementary Table S4.** Descriptive statistics of various physiological traits recorded on 272 rice accessions grown under low and sufficient nitrogen conditions in hydroponics for 25 days.

**Supplementary Table S5**. Rice accessions belonging to N-efficient, N-intermediate, and N-inefficient clusters obtained by hierarchical cluster analysis of phenotypic data.

**Supplementary Table S6**. Mean haplotype trait values of nine contributing traits for *OsNRT1.1A*. Superior and inferior haplogroups were selected based on the mean haplotype trait values.

**Supplementary Table S7**. Mean haplotype trait values of nine contributing traits for *OsNRT1.1B.* Superior and inferior haplogroups were selected based on the mean haplotype trait values.

**Supplementary Table S8**. Mean haplotype trait values of nine contributing traits for *OsNRT1.1C.* Superior and inferior haplogroups were selected based on the mean haplotype trait values.

